# Examining intra-host genetic variation of RSV by short read high-throughput sequencing

**DOI:** 10.1101/2023.05.17.541198

**Authors:** David Henke, Felipe-Andrés Piedra, Vasanthi Avadhanula, Harsha Doddapaneni, Donna M. Muzny, Vipin K. Menon, Kristi L. Hoffman, Matthew C. Ross, Sara J. Javornik Cregeen, Ginger Metcalf, Richard A. Gibbs, Joseph F. Petrosino, Pedro A. Piedra

**Affiliations:** Department of Molecular Virology and Microbiology, Baylor College of Medicine, Houston, TX, USA; Department of Molecular and Human Genetics, Baylor College of Medicine, Houston, TX, USA; Department of Pediatrics, Baylor College of Medicine, Houston, TX, USA

## Abstract

Every viral infection entails an evolving population of viral genomes. High-throughput sequencing technologies can be used to characterize such populations, but to date there are few published examples of such work. In addition, mixed sequencing data are sometimes used to infer properties of infecting genomes without discriminating between genome-derived reads and reads from the much more abundant, in the case of a typical active viral infection, transcripts. Here we apply capture probe-based short read high-throughput sequencing to nasal wash samples taken from a previously described group of adult hematopoietic cell transplant (HCT) recipients naturally infected with respiratory syncytial virus (RSV). We separately analyzed reads from genomes and transcripts for the levels and distribution of genetic variation by calculating per position Shannon entropies. Our analysis reveals a low level of genetic variation within the RSV infections analyzed here, but with interesting differences between genomes and transcripts in 1) average per sample Shannon entropies; 2) the genomic distribution of variation ‘hotspots’; and 3) the genomic distribution of hotspots encoding alternative amino acids. In all, our results suggest the importance of separately analyzing reads from genomes and transcripts when interpreting high-throughput sequencing data for insight into intra-host viral genome replication, expression, and evolution.

## Introduction

A viral infection involves a replicating and therefore evolving population of viruses. The level and distribution of diversity at a given time after infection will depend on the size and composition of the inoculum, the duration of viral replication, how rapidly viral genetic variation is produced de novo, and the nature of host selective pressures (initial screening via secreted antibodies, the innate immune response, and clearing of infected cells by the cellular immune response). Several studies suggest that respiratory viruses like respiratory syncytial virus (RSV) and influenza undergo mostly neutral evolution within a single host during natural infection (1-3), but few report on the expected levels and distribution of genetic variation and it is unknown to what extent different immune functions might constrain viral evolution.

Here we determined the genetic variation contained within intra-host populations of RSV infecting members of a group of previously described adult hematopoietic cell transplant (HCT) recipients (4-7). Cancer patients undergoing myeloablative conditioning require HCT to restore a healthy supply of resident bone marrow cells, including leukocytes such as T and B lymphocytes, neutrophils and macrophages that play essential roles in the host immune response to viral infections. The majority of HCT recipients considered here were fully engrafted at the time of infection and experienced mostly mild disease (4-7).

We sequenced capture probe-derived (Twist Biosciences, Inc.) RSV cDNAs in nasal wash samples from HCT recipients naturally infected with RSV and separately analyzed reads derived from genomes and transcripts, and assessed both data sets for levels and distribution of variation using calculations of per position Shannon entropy. Shannon entropy provides an elegant metric of variation well suited to analyses of high-throughput sequencing data. We found low levels of total genetic variation within the RSV infections studied here, and interesting differences in the levels and distribution of genetic variation contained within genome- and transcript-derived read sets.

## Results

### i. Patient sample data

Nasal wash samples were obtained from a previously described cohort of hematopoietic cell transplant (HCT) recipients naturally infected with respiratory syncytial virus (RSV) (Table 1) and were subjected to short read high-throughput sequencing (NovaSeq Illumina). Patients were infected with either of two widely circulating RSV genotypes (A/Ontario or B/Buenos Aires) and shed virus for either less than 14 days or more (Table 1). Shedding time correlated with transplant type (autologous vs. allogeneic), with patients receiving an autologous HSC transplant tending to show shorter viral shedding times and a more robust neutralizing antibody response (Table 1). A nasal wash sample was collected from each patient at the time of study enrollment and approximately weekly for up to 4 weeks (Table 2). A subset of all samples were successfully sequenced (≥ 90% coverage of whole RSV genome at ≥ 1x sequencing depth) and a further subset were sequenced at a depth permitting downstream analyses to be described (Table 2). Additionally, because of the sequencing methodology employed, it was possible to separately analyze reads from genomes and transcripts.

**Table 1.** Demographics of RSV-infected HCT recipients. A group of previously described hematopoietic cell transplant (HCT) recipients with laboratory-confirmed RSV infection and negative chest radiography findings were identified and enrolled as part of a larger efficacy study within 72 hours of RSV diagnosis (4-7). Patients shed RSV for either less than 14 days or more. Shedding time correlated with transplant type (autologous vs. allogeneic), with patients receiving an autologous HSC transplant tending to show shorter viral shedding times and a greater neutralizing antibody response at convalescence (i.e., 14-60 days after hospitalization).

|  | RSV<br>shedding time<br>(days) |  | p-value* |
| --- | --- | --- | --- |
|  | < 14 | ≥ 14 |  |
| Age (y),<br>mean ± st. dev. | 53.2±14.0 | 46.3±18.5 |  |
| % female (n) | 57.1 (12) | 42.1 (8) |  |
| Race |  |  |  |
| % White (n) | 45.0 (9) | 65.0 (13) |  |
| % Black (n) | 15.0 (3) | 5.0 (1) |  |
| % Hispanic (n) | 30.0 (6) | 30.0 (6) |  |
| % Asian (n) | 10.0 (2) | 0.0 (0) |  |
| Transplant type |  |  |  |
| % autologous (n) | 45.0 (9) | 10.0 (2) | significant |
| % allogeneic (n) | 55.0 (11) | 90.0 (18) | significant |
| Time from HCT<br>(days),<br>median (range) | 169<br>(6-945) | 100<br>(5-1067) |  |
| Acute RSV Nt Ab<br>titer (log2) | 7.0 | 6.2 |  |
| Convalescent RSV<br>Nt Ab titer (log2) | 10.2 | 8.2 | significant |

**Table 2.** Basic sample sequencing information for subgroups of HCT recipients and different read types. HCT recipients were naturally infected with either of two widely circulating RSV genotypes (A/Ontario [A/ON] or B/Buenos Aires [B/BA]) and shed virus for either less than 14 days or more. A nasal wash sample was collected from each patient at the time of study enrollment (i.e., day 0) and approximately weekly for four weeks. A subset of all samples were successfully sequenced at ≥ 90% coverage of whole RSV genome and ≥ 1x sequencing depth; a further subset were sequenced at a depth permitting downstream analyses to be described (≥ 90% coverage of whole RSV genome at ≥ 10x sequencing depth). Additionally, because of the sequencing methodology employed, it was possible to separately analyze reads from genomes and transcripts (Gen and Trx, respectively). **(a)** Basic summary information for samples sequenced at 1x read depth across ≥ 90% of the reference RSV genome (RSV/A/ON or RSV/B/BA). **(b)** Basic summary information for samples sequenced at 10x read depth across ≥ 90% of the reference RSV genome (RSV/A/ON or RSV/B/BA).

**Sequencing: 90% coverage at 1x depth**
| Infecting virus | RSV shedding time (days) | # subjects | Samples per subject, mean [range] | Duration of illness covered (days), mean [range] |
| --- | --- | --- | --- | --- |
| RSV/A/ON | < 14 | 7 | 1.6 [1-2] | 3.4 [0-5] |
|  | ≥ 14 | 9 | 2.7 [2-5] | 12.8 [5-25] |
| RSV/B/BA | < 14 | 8 | 2.0 [1-3] | 2.9 [0-5] |
|  | ≥ 14 | 7 | 3.4 [2-6] | 12.3 [5-28] |

| Infecting virus | RSV shedding time (days) | Read type (Gen/Trx) | # subjects | Samples per subject, mean [range] | Duration of illness covered (days), mean [range] |
| --- | --- | --- | --- | --- | --- |
| RSV/A/ON | < 14 | Gen | 4 | 1.0 [1-1] | 1.7 [0-5] |
|  |  | Trx | 7 | 1.6 [1-2] | 3.0 [0-5] |
|  | ≥ 14 | Gen | 7 | 1.6 [1-4] | 2.6 [0-14] |
|  |  | Trx | 9 | 2.7 [2-5] | 11.2 [5-25] |
| RSV/B/BA | < 14 | Gen | 5 | 1.4 [1-2] | 1.8 [0-4] |
|  |  | Trx | 8 | 2.0 [1-3] | 2.9 [0-5] |
|  | ≥ 14 | Gen | 5 | 1.4 [2-6] | 2.6 [0-11] |
|  |  | Trx | 7 | 3.4 [2-6] | 12.3 [5-28] |

### ii. Varying read depth and variation in sequenced RSV genomes and transcripts

We began our analysis by plotting sequencing or read depth across ON and BA reference genomes for data derived from 1) genomes and 2) transcripts (Fig 1). The latter should also reflect the contribution of low-abundance anti-genomes. All 4 data sets show fairly uniform coverage across the RSV genome (Fig 1), with the average read depth from transcripts exceeding that from genomes by approximately 100-fold.

**Fig 1.**
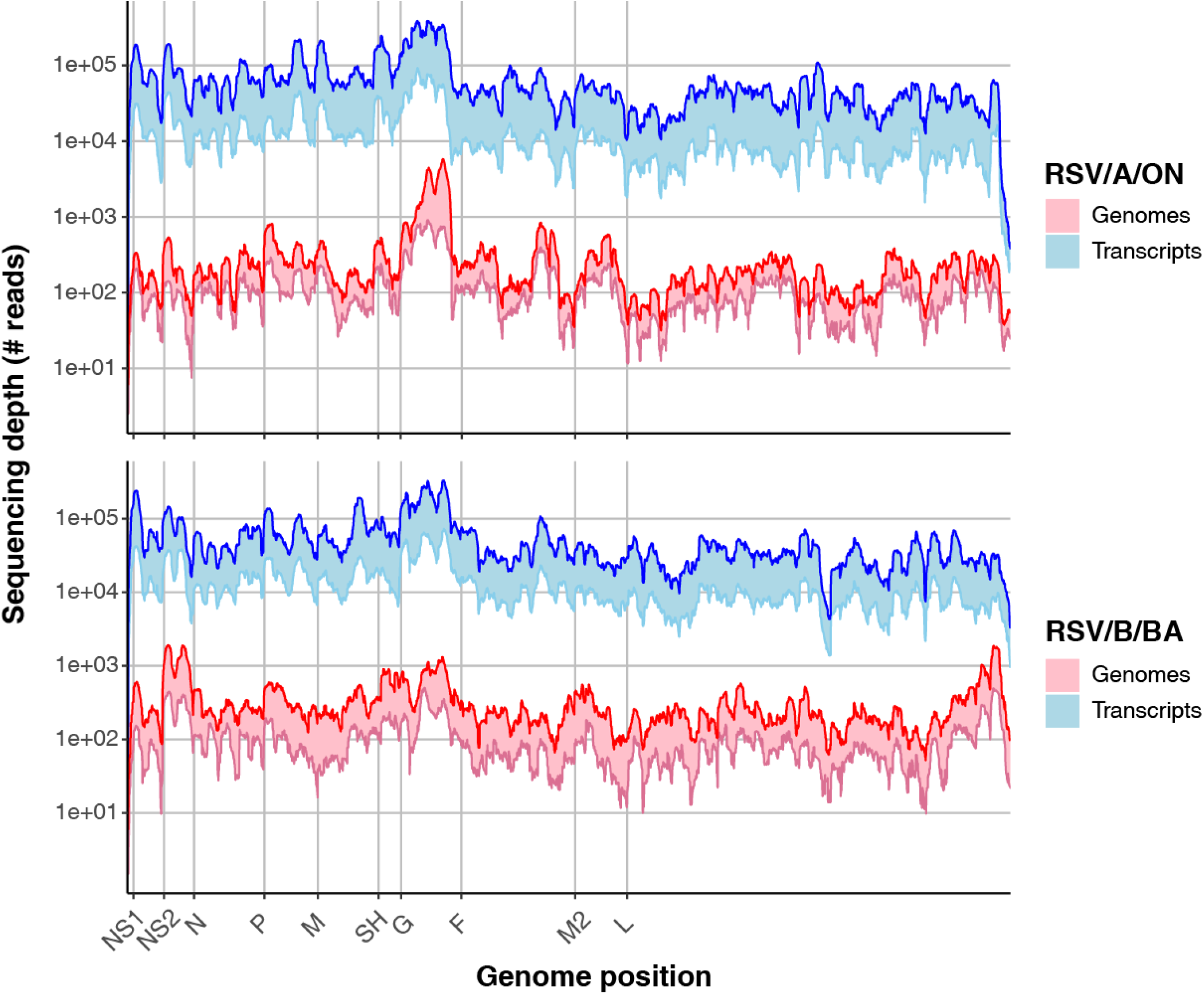
Transcripts exceed genomes by ∼100x and reads derived from both show fairly uniform coverage of reference RSV genomes. The interquartile range of per position sequencing depth from genome-(in red) and transcript-derived read sets (in blue) is plotted along the RSV genome for both RSV/A/ON (top plot) and RSV/B/BA references (bottom plot). Darker lines represent the upper bounds of Q3 and Q1.

In order to begin characterizing the genetic variation supported by the intra-host populations of infecting RSV sequenced here, we adopted an approach based on measuring the Shannon entropy (*H*) of every nucleotide position in our sequencing data set (Fig 2). Plots of per position Shannon entropy across the two reference RSV genomes reveal varying levels of variation across the RSV genome and across samples, with entropy values from genome derived-reads generally exceeding those from transcripts (Fig 2). Calculations of average or bulk Shannon entropy per sample make clear that sequenced RSV genomes show greater variation than sequenced RSV transcripts (Fig 3). Restricting our attention to mean values from day 0 samples, genomes show 4-5x more variation than transcripts (Fig 3), although the bulk Shannon entropy is low across samples. For instance, the maximum per sample average Shannon entropy found (=0.11) would in the simplest case of two possible ‘alleles’ (A or G, say) correspond with a minority ‘allele’ abundance of just over 2%. Thus, whether analyzing reads from RSV genomes or transcripts, the level of genetic variation supported by an infecting population of RSV within a single host is low in the samples tested.

**Fig 2.**
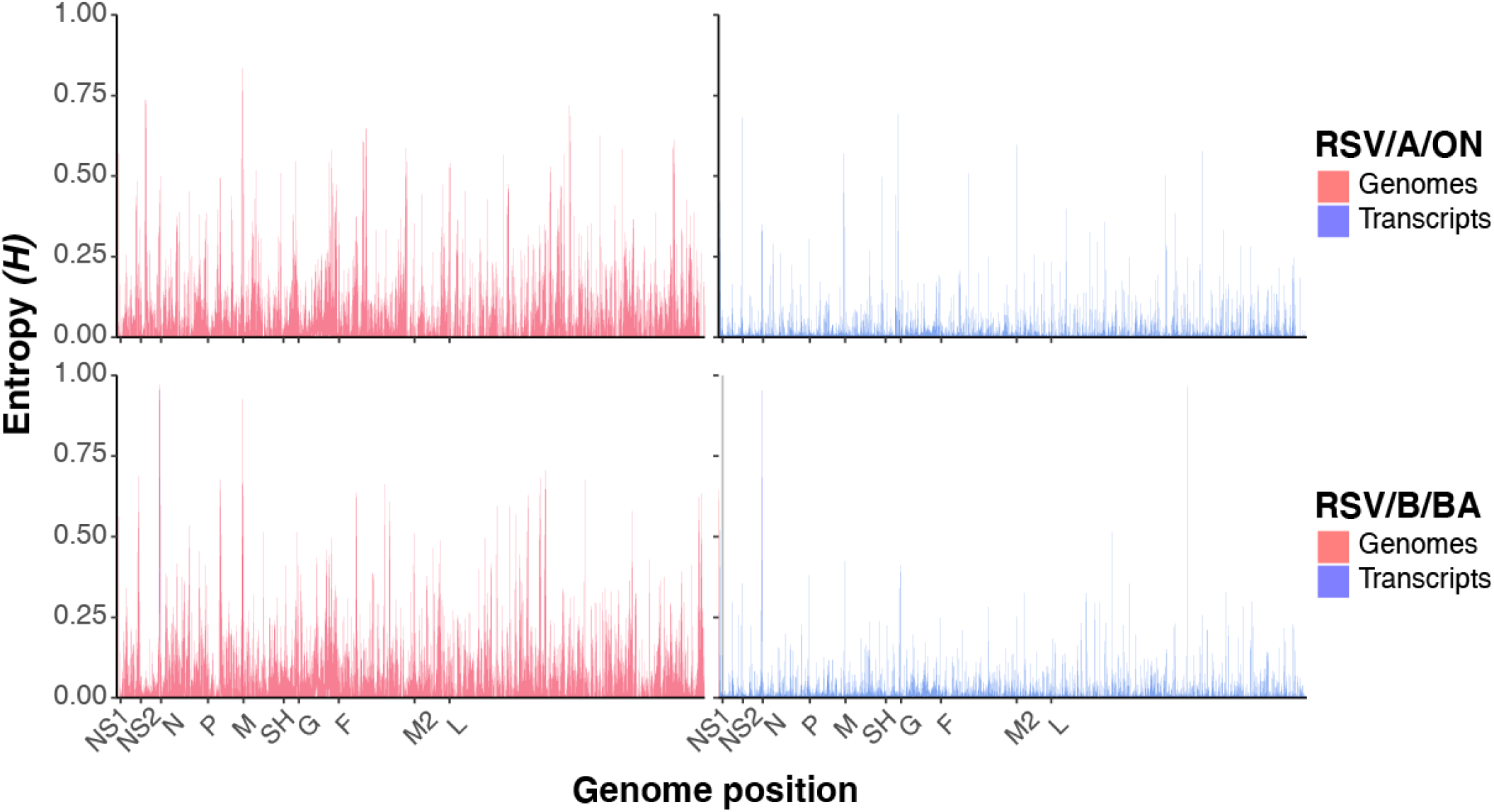
Genomes are more variable than transcripts but both contain highly variable positions located across the RSV genome in different samples. The interquartile range of per position Shannon entropy (*H*) from genome-(in red) and transcript-derived read sets (in blue) is plotted along the RSV genome for both RSV/A/ON (top plot) and RSV/B/BA references (bottom plot).

**Fig 3.**
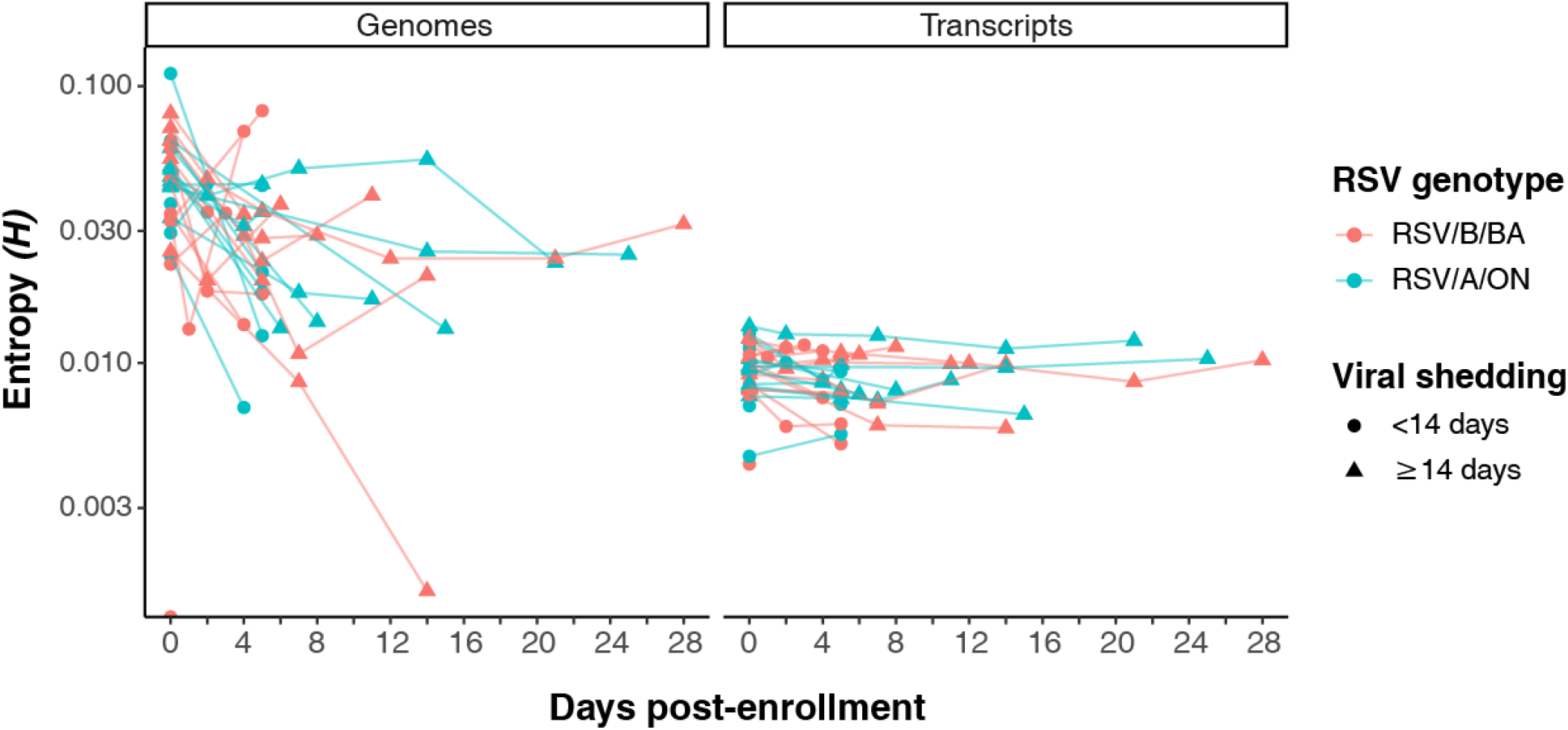
Per sample average or bulk Shannon entropies of genomes and transcripts differ in magnitude and dynamics. Plots of per sample average Shannon entropy (*H*) vs. day of sample acquisition. All per position Shannon entropies for single samples were averaged for genome-(left plot) and transcript-derived read sets (right plot). RSV/A/ON data in blue; RSV/B/BA data in red; data from subjects who shed RSV for < 14 days in closed circular points; data from subjects who shed RSV for ≥ 14 days in closed triangular points.

We also observed that the more variable genomes showed a general drop in bulk entropy over time, while transcript entropies appeared more stable (Fig 3). There are exceptions to the decline in genome entropies over time: one patient showed a bulk entropy maximum at day 14, and a few showed sharp increases (Δ*H* ≈ 0.02) over 2 to 5 days (Fig 3). The former patient shed RSV for longer than 14 days, and most cases showing an increase in bulk genome entropy over any window of time came from longer shedders (Fig 3).

### iii. Distribution of hotspots and estimates of functional variation

Our initial analysis of per position and bulk Shannon entropies from genome- and transcript-derived reads showed low levels of genetic variation within intra-host populations of infecting RSV. However, bulk or average per sample Shannon entropies mask the existence of positions showing exceptionally high variation. Thus, we decided to analyze our data for such ‘hotspots’ (*H* ≥ 0.3) and to determine their distribution across the RSV genome. For this analysis, we restricted our attention to genome- and transcript-derived data sets showing at least 10x coverage across 90% of the reference genome. In both data sets, a minority of positions show a Shannon entropy high enough to be considered hotspots (Fig 4). However, consistent with the differences observed between sequenced RSV genomes and transcripts, RSV genomes are ∼20-fold more enriched for such hotspots (∼3.7% vs. 0.2% of all positions analyzed per sample). In addition, the genomic distribution of hotspots is much more uniform across non-coding and coding sequences in genome-than transcript-derived reads and variation in the latter has a strong tendency to cluster in non-coding sequences (Fig 4).

**Fig 4.**
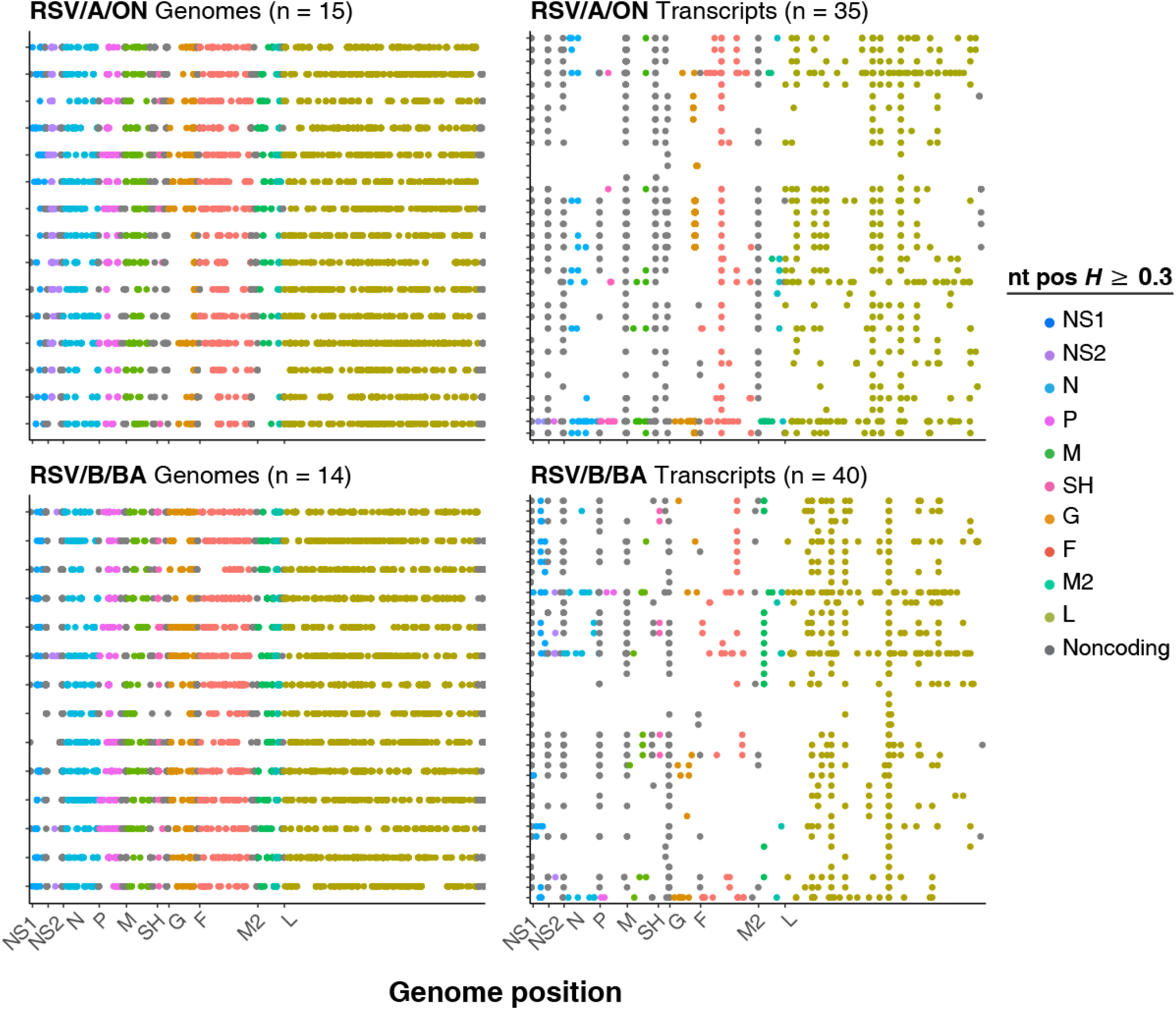
‘Hotspots’ of variation are more numerous and densely distributed across the RSV genome in genome-derived reads than transcripts. Hotspots or single positions showing exceptionally high variation (Shannon entropy (*H*) ≥ 0.3) are plotted across the genome for both genome-(left plots) and transcript-derived read sets (right plots) from RSV/A/ON (top plots) and RSV/B/BA (bottom plots) infections. Each plot contains data for the number of samples indicated in parentheses. Each line in each plot shows the genomic distribution of hotspots for a single sample. Hotspots are colored according to their position within either 1) different noncoding sequences or 2) the 10 coding sequences of RSV.

We further analyzed these hotspots for obvious functional variation by determining whether hotspots within coding sequences encoded alternative amino acids. Interestingly, whether from genomes or transcripts, approximately 50% of all hotspots identified encoded alternative amino acids (Fig 5). The number and distribution of these sites varied from sample to sample, and more highly in transcript-than genome-derived reads (Fig 5).

**Fig 5.**
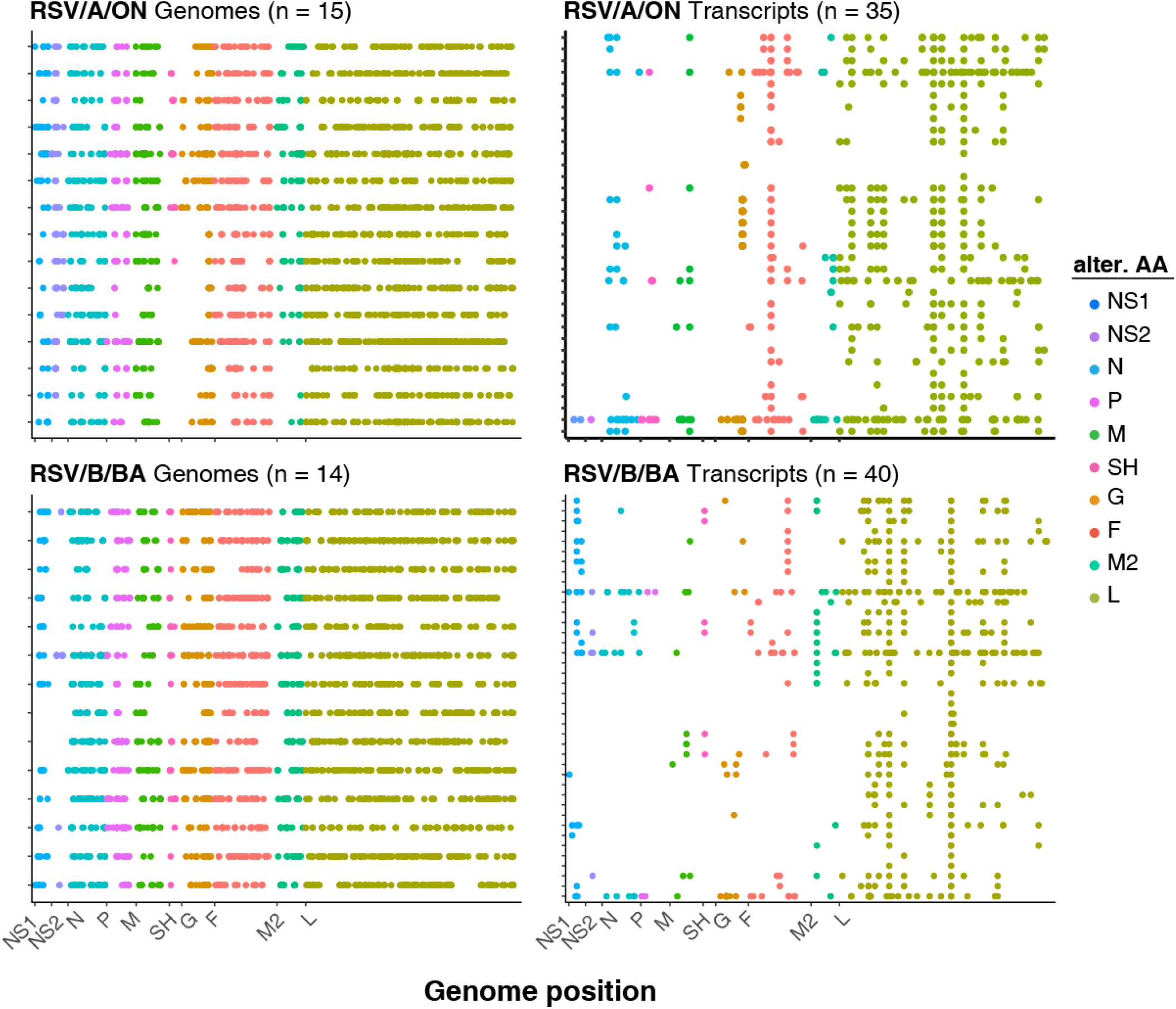
‘Hotspots’ of functional variation are more numerous and densely distributed across the RSV genome in genome-derived reads than transcripts. Hotspots or single positions showing exceptionally high variation (Shannon entropy (*H*) ≥ 0.3) and encoding alternative amino acids are plotted across the genome for both genome-(left plots) and transcript-derived read sets (right plots) from RSV/A/ON (top plots) and RSV/B/BA (bottom plots) infections. Each plot contains data for the number of samples indicated in parentheses. Each line in each plot shows the genomic distribution of hotspots encoding one or more alternative amino acids for a single sample. Hotspots are colored according to their position within the 10 coding sequences of RSV.

Per position Shannon entropies were recalculated for the amino acid (AA) sequences derived from nucleotide hotspots within coding sequences (Fig 6). For both genome- and transcript-derived read sets, the level of variation for any given AA hotspot varies greatly from sample to sample, but genome-derived reads show a much greater number of and more highly distributed AA hotspots than transcript-derived reads (Fig 6).

**Fig 6.**
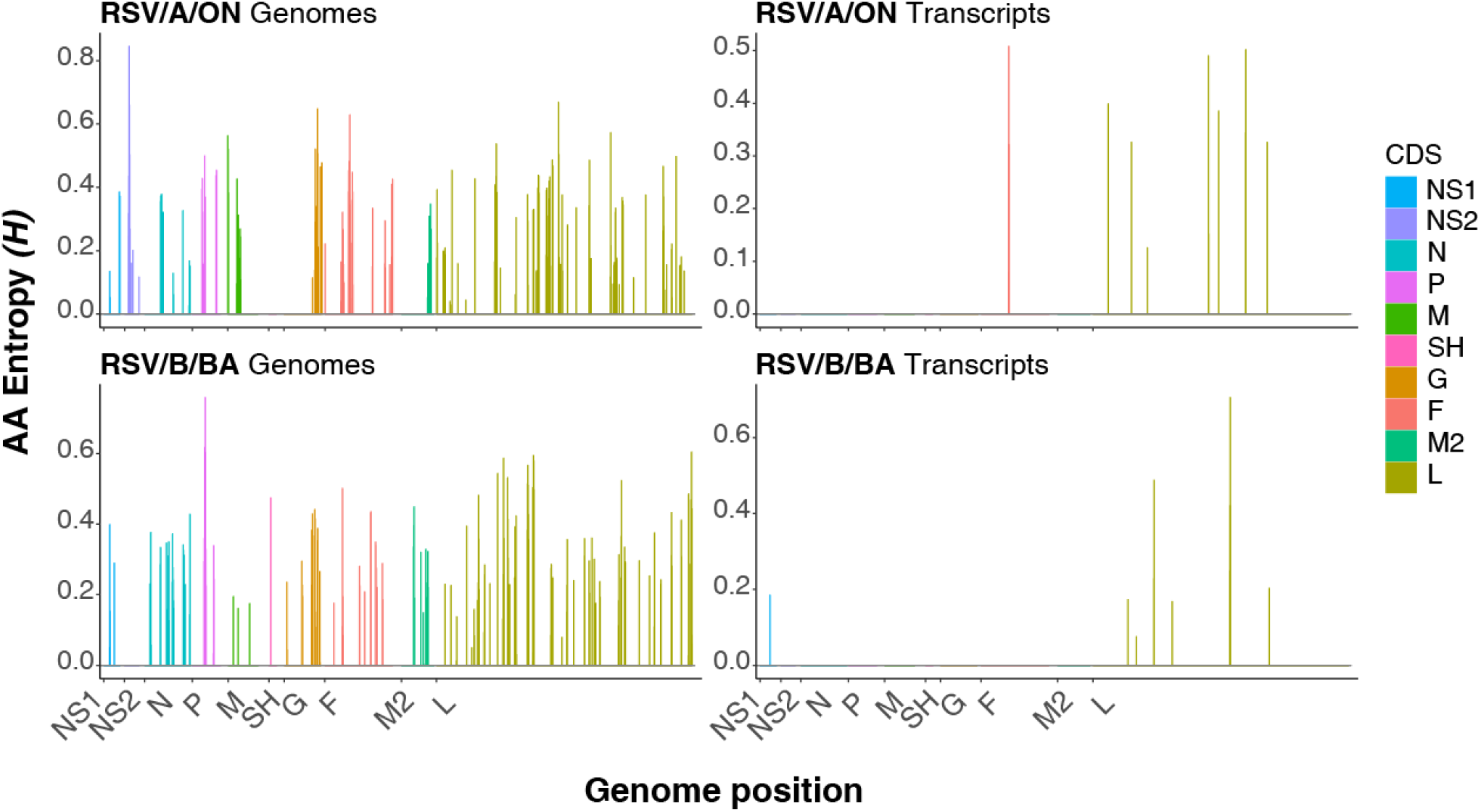
Amino acid (AA) Shannon entropies are higher across the RSV genome in genome-derived reads than transcripts. The interquartile range (IQR) of per position amino acid Shannon entropy (*H*) from genome-(left plots) and transcript-derived read sets (right plots) is plotted along the RSV genome for both RSV/A/ON (top plots) and RSV/B/BA references (bottom plots). Bars showing the IQR of AA *H* are colored according to their position within the 10 coding sequences of RSV.

## Discussion

Our study revealed generally low levels of genetic variation with interesting differences between genome- and transcript-derived read sets from intra-host populations of RSV infecting a cohort of adult HCT recipients.

The 100-fold difference observed in average read levels between genomes and transcripts is consistent with the expectation established from in vitro measurements. These results suggested our sequencing data were minimally biased to different regions of the RSV genome and to either of the two major species of viral nucleic acid present during RSV infection (transcripts and genomes). However, reads mapping to the G gene are slightly more abundant in both genomes and transcripts, especially those derived from RSV/A/ON infections. This appears consistent with multiple studies showing higher than expected levels of the G gene (8-11), especially G gene mRNA. However it may also reflect a subtle sequencing bias of unknown origin, as it is present in both genome- and transcript-derived reads. There is also a noticeable bump in reads mapping to the NS2 gene from RSV/B/BA genomes. This fluctuation appears specific to RSV/B/BA genomes and may reflect a larger proportion of variant genomes (perhaps partly or fully defective viral genomes) containing the NS2 gene along with a subset of the remaining RSV genes. This might also reflect a subtle sequencing bias.

Although both genome- and transcript-derived read sets showed a number of high entropy positions across reference genomes in different samples, the vast majority of positions showed little variation. For example, and as mentioned previously, the largest average or bulk Shannon entropy calculated for a single sample was 0.11, which equals a minority ‘allele’ abundance of just over 2% assuming the simplest case of only two possibilities (A or G, say). The average bulk Shannon entropy for a given sample is closer to 0.03 for genomes and 0.01 for transcripts. The former value corresponds to a minority ‘allele’ abundance of around 0.5% (again assuming only two possible ‘alleles’). Nevertheless, genome sequences clearly contained greater variation than transcripts (approximately 4-5x more) and bulk genome entropies from different patients showed more interesting dynamics, generally dropping over time, while bulk transcript entropies were more stable. This might reflect a purifying selection of viral genomes within the host while the greater stability of the lower transcript entropies may be a consequence of a time-independent error rate for transcribing RSV polymerases. There were exceptions to the decline in genome entropies over time, with most samples showing an increase over any time interval coming from patients who shed RSV for ≥ 14 days (vs. < 14 days). This might reflect, albeit very subtly, the somewhat greater permissiveness of these hosts.

Genome- and transcript-derived reads showed interesting differences in the number and distribution of variation ‘hotspots’ (*H* ≥ 0.3). We chose a Shannon entropy of ≥ 0.3 to identify variation hotspots because *H* = 0.3 corresponds with a rather large minority ‘allele’ abundance of 10% assuming two possibilities (A and G, say). Genomes showed approximately 20-fold more hotspots than transcripts, and the distribution of hotspots was much more uniform across non-coding and coding sequences in genome-than transcript-derived reads. Indeed, variation in the latter had a strong tendency to cluster in non-coding sequences, especially when considering the density of hotspots (i.e., the number of hotspots within a given region divided by the number of positions within that region). Indeed, transcript hotspots appear to be a non-random subset of genome hotspots, potentially indicating the contribution of transcriptionally mute defective viral genomes to our sequencing data (12, 13).

We further analyzed variation hotspots for clear functional variation by determining whether hotspots within coding sequences encoded alternative amino acids. Approximately 50% of all hotspots identified encoded alternative amino acids whether from genomes or transcripts. We thus estimated that the percentage of all coding sequence positions in the RSV genome encoding alternative amino acids was ∼2% from sequenced genomes and ∼0.1% from transcripts within our data. As observed throughout this study, and consistent with the presence of defective viral genomes (12, 13), the variation contained within transcripts is a subset of that contained within genomes.

Here we made use of the ability to separately analyze genome- and transcript-derived reads from high-throughput sequencing data to characterize the levels and distribution of genetic variation contained within natural infections of RSV. Future studies might involve patient populations containing greater differences in host immune status to better search for an immune-mediated effect on the magnitude, distribution, and evolution of viral genetic variation within single infections. It would also be ideal to collect data from a larger number of patients and more densely through time – Grad et al. sequenced 26 samples over more than 2 months from a single infant infected with RSV (14) – while optimizing sample collection for the generation of high-quality sequencing data. Finally, subjecting samples to long read sequencing in order to resolve variant viral genomes including defective viral genomes would be highly informative.

## Methods

### i. Study population

A group of previously described hematopoietic cell transplant (HCT) recipients with laboratory-confirmed RSV infection and negative chest radiography findings were identified from 2012 to 2015 (4-7). Patients were enrolled as part of a ribavirin efficacy trial within 72 hours of RSV diagnosis. Longitudinal nasal wash (NW) samples were collected at enrollment (i.e., day 0), day 2-7, and weekly up to 29 days post-enrollment. The study protocol was approved by the institutional review boards of Baylor College of Medicine and the University of Texas MD Anderson Cancer Center. Written informed consent was obtained from all participants.

### ii. Sample preparation and sequencing

Viral RNA was extracted from NW samples using the Mini Viral RNA Kit (QIAGEN Sciences, Maryland, USA) on the automated platform QIAcube (QIAGEN, Hilden, Germany) according to the manufacturer’s instructions. Pooled cDNA libraries were hybridized with biotin-labeled probes from the RSV Panel (Twist Biosciences, Inc.) at 70°C for 16 hours according to (15). The RSV probe set size was 23.77 Mb and was designed based on 1,570 publicly available genomic sequences of RSV isolates. In this probe set there are 87,025 unique probes of 80 bp length which cover 99.79% of the targeted isolates. Captured virus targets were incubated with streptavidin beads for 30 minutes at room temperature. Streptavidin beads bound with virus targets were washed and amplified with KAPA HiFi HotStart enzyme. The amount of each cDNA library pooled for hybridization and post-capture amplification PCR cycles (12–20) were determined empirically according to the virus Ct values. In general, between 1.8 to 4.0 μg of pre-capture library were used for hybridization with the probes; post-capture libraries were sequenced on an Illumina NovaSeq S4 flow cell to generate 2×150 bp paired-end reads.

### iii. Sequencing data preparation

Cleaned RNA sequence was called using the VirMap pipeline (16). Sample sequences were aligned to custom RSV reference genomes using Bowtie2 (17). Samtool’s mpileup (18) command was used for read pileup creation for each sample. A custom Python (Python Software Foundation, www.python.org) script was utilized to transform the pileup output into a tabular form.

### iv. Analysis of sequencing data

The sequencing data from each sample was separated into two subsets, genomic and transcriptomic. Unless otherwise noted, a minimum sequencing depth of 10 reads was required at each position across 90% of the reference genome (RSV/A/ON or RSV/B/BA) for each sample to be used in downstream analyses.

### v. Viral sequence Shannon entropy calculations

Shannon entropy (*H*) was defined within each sample and at every genomic position as:

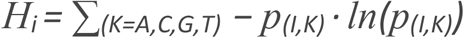

i = sample identified; has a dimension of rows = # genomic positions with coverage ≥ 10 reads and columns = 1

*p* = proportion of base = the number of counts for a given base divided by the total counts at a given genomic position

I = genomic position

A, C, G, and T= nucleotide base type

Analyses were conducted in R 3.4.4 (R Foundation for Statistical Computing, Vienna, Austria) unless otherwise stated.

### vi. Detecting non-synonymous changes and calculating amino acid Shannon entropies

To predict the amino acid (AA) representation across coding sequences from genome- and transcript-derived reads, we assumed a uniform sequencing error rate of nt substitution and only analyzed nt composition (i.e., did not consider insertions/deletions or associated frameshift mutations). Nucleotide counts, binned as the four possible nt bases (A, C, G, T), were determined at each coding position within each sample. The AA abundance was calculated at each position within a codon; if neighboring positions within a codon showed more than one nt base, the majority base(s) was used to determine the AA assignment.

## Acknowledgments

Funding:This work was supported by the NIH Texas Medical Center Genomic Center for Infectious Diseases (TMC-GCID, grant# U19AI144297).

## References

1. Debbink K, McCrone JT, Petrie JG, Truscon R, Johnson E, Mantlo EK, Monto AS, Lauring AS. 2017. Vaccination has minimal impact on the intrahost diversity of H3N2 influenza viruses. PLoS Pathog 13:e1006194.

2. McCrone JT, Woods RJ, Martin ET, Malosh RE, Monto AS, Lauring AS. 2018. Stochastic processes constrain the within and between host evolution of influenza virus. Elife 7.

3. Nelson MI, Simonsen L, Viboud C, Miller MA, Taylor J, George KS, Griesemer SB, Ghedin E, Sengamalay NA, Spiro DJ, Volkov I, Grenfell BT, Lipman DJ, Taubenberger JK, Holmes EC. 2006. Stochastic processes are key determinants of short-term evolution in influenza a virus. PLoS Pathog 2:e125.

4. Avadhanula V, Chemaly RF, Shah DP, Ghantoji SS, Azzi JM, Aideyan LO, Mei M, Piedra PA. 2015. Infection with novel respiratory syncytial virus genotype Ontario (ON1) in adult hematopoietic cell transplant recipients, Texas, 2011-2013. J Infect Dis 211:582–589.

5. Ye X, Cabral de Rezende W, Iwuchukwu OP, Avadhanula V, Ferlic-Stark LL, Patel KD, Piedra FA, Shah DP, Chemaly RF, Piedra PA. 2020. Antibody Response to the Furin Cleavable Twenty-Seven Amino Acid Peptide (p27) of the Fusion Protein in Respiratory Syncytial Virus (RSV) Infected Adult Hematopoietic Cell Transplant (HCT) Recipients. Vaccines (Basel) 8.

6. Ye X, Iwuchukwu OP, Avadhanula V, Aideyan LO, McBride TJ, Ferlic-Stark LL, Patel KD, Piedra FA, Shah DP, Chemaly RF, Piedra PA. 2018. Comparison of Palivizumab-Like Antibody Binding to Different Conformations of the RSV F Protein in RSV-Infected Adult Hematopoietic Cell Transplant Recipients. J Infect Dis 217:1247–1256.

7. Ye X, Iwuchukwu OP, Avadhanula V, Aideyan LO, McBride TJ, Henke DM, Patel KD, Piedra FA, Angelo LS, Shah DP, Chemaly RF, Piedra PA. 2021. Humoral and Mucosal Antibody Response to RSV Structural Proteins in RSV-Infected Adult Hematopoietic Cell Transplant (HCT) Recipients. Viruses 13.

8. Aljabr W, Touzelet O, Pollakis G, Wu W, Munday DC, Hughes M, Hertz-Fowler C, Kenny J, Fearns R, Barr JN, Matthews DA, Hiscox JA. 2016. Investigating the Influence of Ribavirin on Human Respiratory Syncytial Virus RNA Synthesis by Using a High-Resolution Transcriptome Sequencing Approach. J Virol 90:4876–4888.

9. Krempl C, Murphy BR, Collins PL. 2002. Recombinant respiratory syncytial virus with the G and F genes shifted to the promoter-proximal positions. J Virol 76:11931–11942.

10. Piedra FA, Qiu X, Teng MN, Avadhanula V, Machado AA, Kim DK, Hixson J, Bahl J, Piedra PA. 2020. Non-gradient and genotype-dependent patterns of RSV gene expression. PLoS One 15:e0227558.

11. Rajan A, Piedra FA, Aideyan L, McBride T, Robertson M, Johnson HL, Aloisio GM, Henke D, Coarfa C, Stossi F, Menon VK, Doddapaneni H, Muzny DM, Javornik Cregeen SJ, Hoffman KL, Petrosino J, Gibbs RA, Avadhanula V, Piedra PA. 2022. Multiple Respiratory Syncytial Virus (RSV) Strains Infecting HEp-2 and A549 Cells Reveal Cell Line-Dependent Differences in Resistance to RSV Infection. J Virol 96:e0190421.

12. Lopez CB. 2014. Defective viral genomes: critical danger signals of viral infections. J Virol 88:8720–8723.

13. Pathak KB, Nagy PD. 2009. Defective Interfering RNAs: Foes of Viruses and Friends of Virologists. Viruses 1:895–919.

14. Grad YH, Newman R, Zody M, Yang X, Murphy R, Qu J, Malboeuf CM, Levin JZ, Lipsitch M, DeVincenzo J. 2014. Within-host whole-genome deep sequencing and diversity analysis of human respiratory syncytial virus infection reveals dynamics of genomic diversity in the absence and presence of immune pressure. J Virol 88:7286–7293.

15. Doddapaneni H, Cregeen SJ, Sucgang R, Meng Q, Qin X, Avadhanula V, Chao H, Menon V, Nicholson E, Henke D, Piedra FA, Rajan A, Momin Z, Kottapalli K, Hoffman KL, Sedlazeck FJ, Metcalf G, Piedra PA, Muzny DM, Petrosino JF, Gibbs RA. 2021. Oligonucleotide capture sequencing of the SARS-CoV-2 genome and subgenomic fragments from COVID-19 individuals. PLoS One 16:e0244468.

16. Ajami NJ, Wong MC, Ross MC, Lloyd RE, Petrosino JF. 2018. Maximal viral information recovery from sequence data using VirMAP. Nat Commun 9:3205.

17. Langmead B, Salzberg SL. 2012. Fast gapped-read alignment with Bowtie 2. Nat Methods 9:357–359.

18. Li H, Handsaker B, Wysoker A, Fennell T, Ruan J, Homer N, Marth G, Abecasis G, Durbin R, Genome Project Data Processing S. 2009. The Sequence Alignment/Map format and SAMtools. Bioinformatics 25:2078–2079.

